# Therapeutic potential of TAF1 bromodomains for cancer treatment

**DOI:** 10.1101/394254

**Authors:** Veronica Garcia-Carpizo, Sergio Ruiz-Llorente, Jacinto Sarmentero, Maria J. Barrero

## Abstract

The discovery of the antiproliferative effects of BRD4 bromodomain inhibitors prompted us to investigate additional bromodomains that might be involved in supporting cellular proliferation. TAF1 is a general transcription factor with two bromodomains that is likely to play important roles in cell viability by supporting transcription. Our work shows that knock down of TAF1 caused antiproliferative effects in several cancer cell lines. Using CRISPR-Cas9 editing techniques we demonstrate that the bromodomains of TAF1 are essential to maintain proliferation of K562 and H322 cells. BAY-299, the best TAF1 bromodomain inhibitor developed so far, also showed antiproliferative effects. BAY-299 caused discrete transcriptional changes that were likely on target but did not correlate with strong effects in cell cycle distribution suggesting that these effects might be, at least in part, mediated by another target. Compared to the TAF1 knock down, BAY-299 specifically downregulated the expression of genes with high levels of TAF1 and histone acetylation at their promoters, suggesting that targeting the bromodomains has distinct transcriptional effects than targeting the whole protein. This might help to provide a therapeutic window to target specifically cancer cells with high levels of histone acetylation at proliferation-related genes.

## INTRODUCTION

TAF1 (TATA box binding protein-associated factor 1, also known as TAF250) is the largest subunit and core scaffold of the TFIID complex. TFIID binds the core promoter to properly position RNA polymerase II, and serves as a scaffold for the assembly of the pre-initiation complex at the first step of transcription [1]. TFIID is composed of the TATA-binding protein (TBP) and a group of evolutionarily conserved proteins known as TBP-associated factors or TAFs. TAFs may participate in basal transcription, serve as coactivators, function in promoter recognition or modify general transcription factors to facilitate complex assembly and transcription initiation. TAF1 is the largest subunit of TFIID, which interacts with TBP through an N-terminal TBP-binding sequence. TAF1 also binds to core promoter sequences, activators and other transcriptional regulators, and these interactions affect the rate of transcription initiation. TAF1 has been reported to possess various biochemical activities including protein phosphorylation, histone acetylation, and acetylated histone tail recognition activities. These activities have been mapped to the two terminal kinase domains, a central histone acetyltransferase domain, and two tandem bromodomains, respectively.

TAF1 is an interesting therapeutic candidate involved in transcription activation. In addition, TAF1 has been described to be involved in G1 progression through phosphorylation of p53 at Thr-55 resulting in its degradation [2]. During recent years a number of inhibitors able to block the interaction of bromodomains with acetylated histones have been developed, most remarkable inhibitors of the bromodomain of BRD4 able to block the proliferation of cancer cells [3]. However, the outcomes of targeting most of the 60 different bromodomains found in the human proteome remain unknown. Therefore, we explored the therapeutic potential of targeting TAF1 bromodomains. Despite TAF1 being essential for maintaining proper transcription in all cells we speculated that cancer cells might be particularly sensitive to inhibitors of the TAF1 bromodomains by blocking the expression of oncogenes with very high levels of acetylation. Recently, two different types of TAF1 bromodomain inhibitors have been developed [4,5]. Although these were elegant studies the therapeutic potential of targeting TAF1 still remains poorly understood. Here, we use genetic and chemical approaches to better understand the molecular consequences of targeting TAF1 and its bromodomains in cancer cell lines.

## RESULTS

We set up to interrogate the potential involvement of TAF1 bromodomains in the proliferation of cancer cell lines. We first infected the myeloma cancer cell line KMS11 with lentiviruses driving the expression of four different shRNAs against TAF1(Suplementary Figure 1A) and a non-target shRNA in a doxycycline inducible fashion (Supplementary Figure 1B). shRNAs targeting TAF1 caused a decrease in the proliferation of KMS11 cells (Supplementary Figure 1C), most conspicuily sh_579 that caused the most significant depletion of TAF1 and most dramatic effects in proliferation. Three of the shRNAs were also tested for effects in proliferation in the CML cell line K562 (Figure 1). shRNAs sh_579 and sh_670 reduced the levels of TAF1 in the presence of doxycycline (Figure 1A) coincident with effects in the proliferation (Figure 1B). In this cell line, sh_581 showed low efficacy and marginal effects on proliferation. Next, we asked if cancer cell lines representative of solid tumors might be also proliferation sensitive to the TAF1 knock down. Supplementary Figures 2, 3 and 4 show that lung cancer cell lines H1568, H322 and H2291 are proliferation sensitive to TAF1 depletion. In all cases, shRNAs sh_579 and sh_670 were the most efficient shRNAs (Supplementary Figures 2A, 3A and 4A) and correlated with effects in proliferation both in growth experiments in which cells were counted (Supplementary Figures 2B, 3B and 4B) or stained with crystal violet (Supplementary Figures 2C, 3C and 4C).

**Figure 1.**
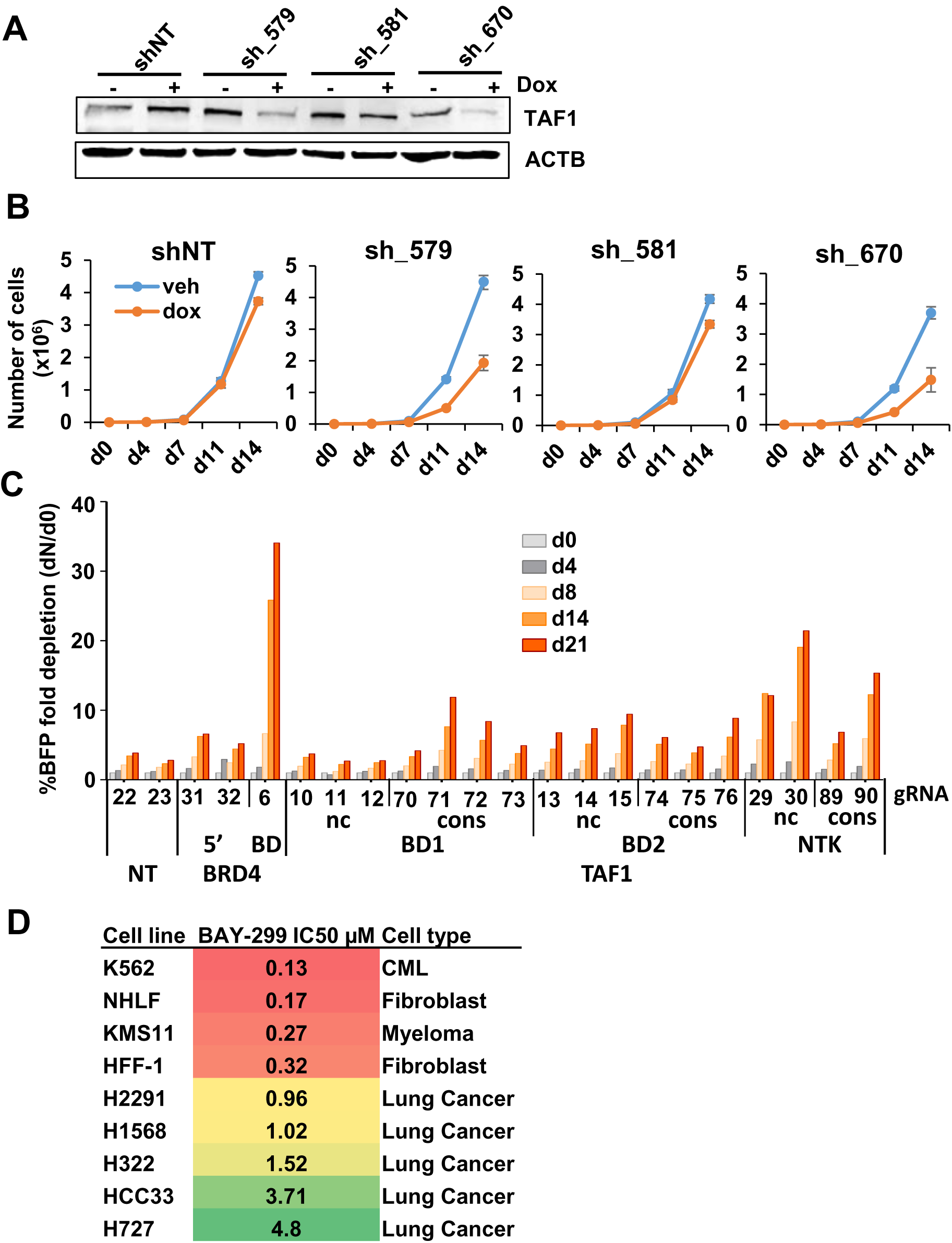
Targeting TAF1 in K562 cells affects proliferation. (a) Western-blot showing the levels of TAF1 in K562 cells transduced with the indicated shRNAs and treated with doxycycline for 7 days. (b) Proliferation curves of the transduced cells treated with vehicle or doxycycline for the indicated days. (c) Growth competition assays in K562 cells transduced with gRNAs targeting diverse domains of TAF1 including conserved (cons) and non-conserved (nc) regions of bromodomains 1 (BD1), bromodomain 2 and N-terminal kinase (NTK) domain. Non-target (NT) gRNAs and gRNAs against BRD4 including 5’ coding region (5’) and bromodomain (BD) were also included as controls. (d) IC50s for BAY-299 in different normal and cancer cell lines.

To confirm the relevance of the TAF1 bromodomains for the proliferation of K562 and H322 cells we used a recently described CRISPR-Cas9 genome editing approach used to evaluate the relevance of protein domains in proliferation [6,7]. This method is based in the fact that one third of randomly introduced mutations are in frame and are likely to generate a full-length protein with mutations in the particular domain targeted by the gRNA. If a domain is relevant for proliferation more pronounced antiproliferative effects will be observed when targeting that domain than an irrelevant domain. We interrogated the effect of introducing mutations in several domains of TAF1 including the two bromodomains and the NTK region. Both conserved and non-conserved amino acids in these regions were targeted by several different gRNAs. We also used gRNAs non-target and targeting the 5’ coding region and the bromodomain one of BRD4 as controls. Figure 1C shows growth competition assays of K562 cells transduced with lentiviruses encoding the indicated gRNAs (BFP+) and non-transduced cells. Targeting the bromodomains or the NTK domain of TAF1 had effects in proliferation that were more dramatic when targeting conserved amino acids (Supplementary Figure 5). Similar results were found in the lung cancer cell lines H322 (Supplementary Figure 3D and Supplementary Figure 6). Therefore, our CRISPR-Cas9 approach suggests that the bromodomains of TAF1 are relevant to sustain the proliferation of the tested cancer cell lines.

As a third approach to evaluate the involvement of TAF1 in proliferation we tested the sensitivity of several cell lines, including the ones used for the knock down, to the recently developed TAF1 bromodomain inhibitor BAY-299 [5]. BAY-299 is the most potent and selective TAF1 bromodomain inhibitor described so far, with an IC50 of 8 nM for the second bromodomain of TAF1. In addition, it also targets with lower potency BRPF2 (IC50 of 67 nM) and TAF1L (IC50 106 nM). Importantly, no significant binding activity towards BRD4 has been reported. Figure 1D shows that the tested cell lines had different sensitivities to BAY-299 being the hematologic cancer cell lines and primary lines of normal human fibroblasts the most sensitive cell lines.

Our next goal was to gain mechanistic insights into the mechanisms of sensitivity of the cell lines to TAF1 depletion and BAY-299 treatment. For that, we first analyzed and compared the transcriptional programs affected by BAY-299 and the two different short hairpins against TAF1 (sh579 and sh670) in K562 (Figure 2). In addition, we compared the responses to the transcriptional consequences of treating K562 cells with the BRD4 bromodomain inhibitor JQ1 and the CREBBP/EP300 bromodomain inhibitor CBP30. Importantly, K562 cells were found proliferation sensitive to the three tested compounds with IC50s of 0.012 µM for JQ1, 0.92 µM for CBP30 and 0.13 µM for BAY-299. Figure 2A shows the amplitude of the transcriptional changes caused by the compounds treatment or the shRNAs. Although bromodomain inhibitors are expected to mainly downregulate gene expression due to interference with the acetylated signal both upregulated and downregulated genes were found after treatments (Figure 2A and 2B). sh579 shRNA caused more discrete transcriptional responses than sh_670, which correlates with the lower efficiency of sh_579 to knock down TAF1 compared to sh_670 (70% and 86% respectively, determined by western blot). Unexpectedly, BAY-299 caused more discrete transcriptional responses than CBP30 (Figure 2A and 2B) even though the sensitivity of K562 cells to BAY-299 was about one order of magnitude higher than to CBP30. The lack of correlation between the amplitude of transcriptional responses and the sensitivity in proliferation to BAY-299 suggests that BAY-299 might be partially mediating its antiproliferative effects through non-transcriptional mechanisms. There was a significant overlap of genes differentially regulated by the knock down and BAY-299, however the overlap between the two shRNAs was more significant than the overlap between each shRNA and BAY-299 (Figure 2C). These discrepancies could be explained by differences in depleting the whole protein versus targeting the bromodomain. In addition, sh_670 shared more downregulated genes with CBP30 and JQ1 than BAY-299 (Figure 2D).

**Figure 2.**
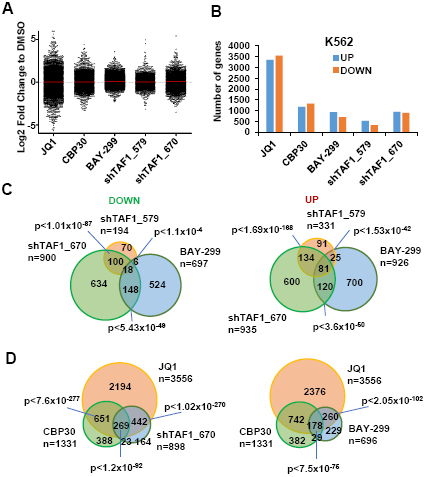
Comparison of transcriptional responses to TAF1 knock down and BAY-299 treatment in K562. (a) Changes in the expression of all genes after TAF1 knock down compared to non-target shRNA or treatment with the indicated compounds compared to DMSO (b) Number of upregulated and downregulated genes at a FDR<0.05. Transcriptional effects after targeting TAF1 in K562 (c) Overlap of upregulated and downregulated genes at a FDR>0.05. (d) Overlap of genes downregulated by the TAF1 knock down (sh_670) or BAY-299 treatment and genes downregulated by CBP30 or JQ1 at a FDR<0.05.P-values for the statistical significance of the overlap between two groups are shown.

We next investigated which transcriptional pathways could be affected by the TAF1 knock down and BAY-299 treatment. Our previous work showed that CBP30 downregulates both the GATA1 and MYC-dependent transcriptional signature in K562 [8]. Like CBP30, sh670 and BAY-299 downregulated genes typically upregulated by MYC and targets of GATA1 in K562 (Figure 3A). In addition, we investigated the enrichment of particular genomic features in genes upregulated or downregulated by the knock down and BAY-299 (Figure 3B). Sh579 showed poor enrichments probably due to the lower efficacy of the knock down. Both sh670 and BAY-299 caused downregulation of a significant number of genes bound by top levels of TAF1, RNA polymerase II (PolII), TBP and H3K27ac around their transcription start sites (TSS) in K562. These enrichments were not found in upregulated genes. Interestingly, both the knock down and BAY-299 affected the expression of genes bound by top levels of TAF1 but only the compound affected those occupied by TAF1 that have very high levels of acetylation around their TSS, suggesting that BAY-299 targets the bromodomains of TAF1. Among these genes we found several histone genes and genes involved in protein translation and metabolic process, all essential to facilitate cell cycle progression (Figure 3C). All these data suggest that BAY-299 causes discrete transcriptional changes that are likely to be mediated through the inhibition of the bromodomains of TAF1, however these do not correlate with the potent antiproliferative effects of this compound.

**Figure 3.**
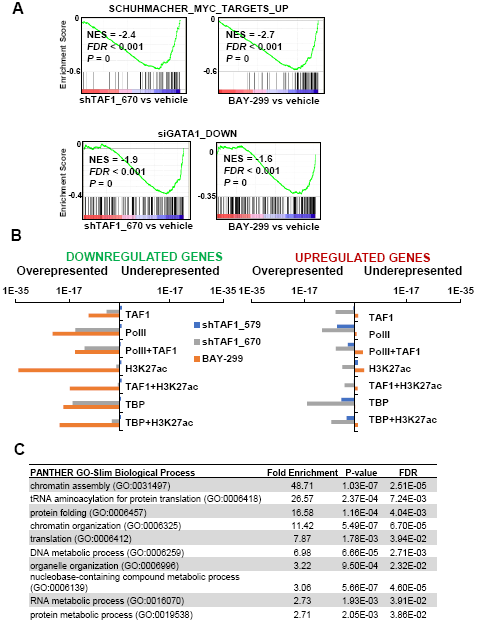
Transcriptional pathways affected by the TAF1 knock down and BAY-299 treatment. (a) GSEA plots showing enrichment of the changes in gene expression caused by TAF1 knock down or BAY-299 treatment for the indicated gene sets. (b) Enrichment of genes upregulated or downregulated by the two shRNAs against TAF1 or BAY-299 in genes occupied by top levels of the indicated factors or histone modifications. (c) PANTHER enrichment in biological processes in genes downregulated by BAY-299 and with top levels of TAF1 and H3K27ac on their TSS.

In order to understand better the differences between the TAF1 knock down and the bromodomain inhibitor we evaluated changes in cell cycle (Figure 4). First, we focused in the changes in cell cycle distribution caused by BAY-299 and its negative control BAY-299N treatments in K562 at 48 and 72 hours of treatment (Figure 4A and 4B). We used as positive control JQ1 that, as previously described [9], caused an increase in the percentage of cells in G0/1 (Figure 4A and 4B). BAY-299 however caused a pronounced increased in the percentage of cells in G2/M in accordance to the sensitivity of K562 cells to this compound. The negative control compound BAY-299N did not have significant effects in cell cycle distribution. Similar results were found in the lung cancer cell line H322 (Supplementary Figure 7A and 7B). We next quantified effects on cell cycle caused by the TAF1 knock down (Figure 4C). Contrary to the compound effect, the TAF1 knock down caused more discrete although significant increases in the number of cells in G0/1 in K562, more significantly for the sh670. Similar results were found in H322 cells (Supplementary Figure 7C). Combinations of knock down and compound treatment (Figure 5A and B) showed that the TAF1 knock down (sh670) only partially rescues the effects of BAY-299 suggesting that this compound might be mediating effects in the cell cycle independently of TAF1. It is interesting to notice that JQ1 and the TAF1 knock down showed synergistic effects on the arrest of cells in G0/1 (Figure 5A and 5B). This result is in agreement with recent data reporting a cross talk between BRD4 and TAF1 [10].

**Figure 4.**
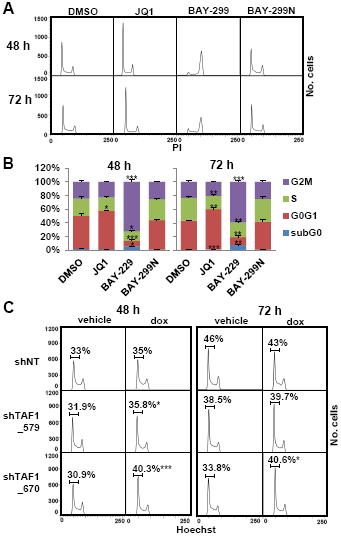
Changes in cell cycle caused by BAY-299 treatment and TAF1 knock down in K562. (a) Cell cycle distribution of K562 cells treated with 200nM JQ1 or 2µM BAY-299 or BAY-299N the indicated compounds for 48 and 72 hours (b) Percentage of cells in each phase of the cell cycle. Mean and standard deviation from three replicates are shown. (d) Cell cycle distribution of K562 cells treated with vehicle or doxycycline for 5 days, reseeded at low confluency and harvested at 48 and 72 hours. The mean from three replicates of the percentage of cells in G0/G1 is indicated. *p<0.05, **p<0.005, ***p<0.0005

**Figure 5.**
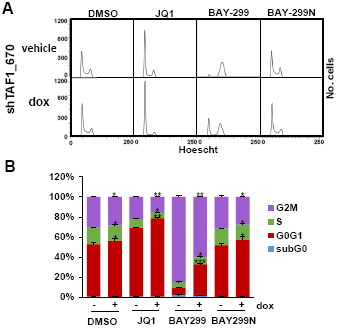
Changes in cell cycle caused by combinations of TAF1 knock down and BAY-299 in K562. (a) Cell cycle distribution of cells pretreated with doxycycline for 5 days, seeded at low confluency and treated with the compounds for 48 hours. (b) Percentage of cells in each phase of the cell cycle. Mean and standard deviation from three replicates are shown. P-values for comparisons between vehicle and doxycycline are shown *p<0.05, **p<0.005, ***p<0.0005.

## DISCUSSION

The discovery of the antiproliferative effects of BRD4 bromodomain inhibitors prompted us to study of additional bromodomains that might be involved in supporting cellular proliferation. BRD4 bromodomain inhibitors prevent the interaction of BRD4 to acetylated histones and block its recruitment to promoters to boost transcription. These inhibitors have been described to affect the expression of oncogenes with high levels of acetylation on their regulatory regions affecting in this way the proliferation of cancer cells. TAF1 is a bromodomain containing protein involved in transcriptional activation that might offer similar therapeutic opportunities to BRD4. However, the therapeutic potential of targeting TAF1 has only been weakly explored mainly due to its essentiality for cell viability as a general transcription factor. We speculated that targeting the bromodomains of TAF1 could offer a therapeutic window that might render cancer cells with high levels of histone acetylation at critical oncogenes more sensitive to inhibition than normal cells. However, the relevance of TAF1 bromodomains for TAF1 function and for transcriptional activation in general was practically unknow at the time we started this study.

Our results show that as expected, depletion of TAF1 compromises the viability of the tested cell lines. Through a CRISPR-Cas9 based gene editing technique we have confirmed that the bromodomains of TAF1 are relevant for the proliferation of cancer cells. To further confirm this fact, we have tested the best TAF1 bromodomain inhibitor described so far, BAY-299. Surprisingly, BAY-299 had strong effects in K562 proliferation that did not correlate with rather discrete transcriptional responses, suggesting that BAY-299 might be, at least partially, mediating antiproliferative effects independent of its effects in transcription. However, several facts suggest that the transcriptional effects mediated by BAY-299 are on target. First, we found a significant overlap of genes downregulated by the TAF1 knock down and BAY-299 treatment, and both the knock down and BAY-299 affected GATA1 and MYC transcriptional programs in K562. Second, there was a significant overlap of genes downregulated by JQ1 or CBP30 and BAY-299, as expected from treatments that affect the readout of acetylation. Finally, genes downregulated by BAY-299 are occupied by TAF1 and have high levels of acetylation in K562. Despite this, analysis of the cell cycle showed that BAY-299 causes an increase in the number of cells in G2/M while the TAF1 knock down causes a discrete but significant increase in the percentage of cells in G0/1. Importantly, JQ1 also causes an increase in the percentage of cells in G0/1 that correlates with the overlap in transcriptional changes caused by JQ1 and the TAF1 knock down. Importantly, we have previously described that CBP30 also increases the number of cells in G0/1 in K562 [8] which also correlates with the overlap in transcriptional responses to CBP30 and TAF1 knock down. In summary, there was a correlation between the transcriptional responses caused by JQ1, CBP30 and the TAF1 knock down and effects on the cell cycle while BAY-299 had completely different effects on the cell cycle despite having transcriptional effects that overlapped with JQ1, CBP30 and the TAF1 knock down. These results suggest that BAY-299 causes transcriptional responses that are on target but effects in the cell cycle that are likely independent of TAF1 inhibition. The fact that BAY-299 still causes an increase the percentage of cells in G2/M in lines depleted of TAF1 further confirms this idea.

Importantly, the analysis of enrichment in genomic features revealed that even though BAY-299 and the TAF1 knock down downregulated the expression of genes occupied by TAF1, BAY-299 specifically downregulated the expression of genes that in addition have very high levels of acetylation at their promoters. This suggests that targeting the bromodomains is likely to have distinct transcriptional effects than targeting the whole protein. Weather this might help to provide a therapeutic window to target specifically cancer cells remains to be determined. In addition, our CRISPR-Cas9 approach also identified the NTK as a potential therapeutic domain.

We conclude that TAF1 bromodomains hold therapeutic potential but compound BAY-299, which is not fully characterized, is probably causing antiproliferative effects at least in part by inhibiting other targets. More selective inhibitors of the TAF1 bromodomains will need to be developed to further evaluate this target.

## MATERIALS AND METHODS

### Cell lines and reagents

Human cancer cell lines K562, H322, H1568, HCC33, H727, HFF-1 and H2291 were purchased from ATCC. NHLF were purchased from LONZA. KMS11 was purchased from JCRB. The identity of the cell lines was verified with STR analysis. qPCR-based mycoplasma test was routinely carried out once a month. Antibodies used for western-blot were TAF1 sc-735 from Santa Cruz and ACTB (Ac-15) A5441 from Sigma Aldrich. BAY-299 and CBP30 were purchased from Tocris Bioscience. BAY-299N was purchased from Sigma-Aldrich. JQ1 was purchased from Selleck Chemicals.

### Proliferation assays

For proliferation curves cells were counted using a haemocytometer and plated at day 0 in triplicate for each condition, treated with 0.5 µ g/ml doxycycline and harvested and counted at the indicated days. For colony forming assays cells were plated in triplicate and stained with crystal violet at the indicated day after plating. Crystal violet staining was quantified using ImageJ. To determine IC50s cells were grown in 96-well plates in the presence of increasing amounts of compound. At day seven viability was determined using the CellTiter-Glow Luminescent Assay. IC50 values were calculated with a four-parameter variable-slope dose response curve using the GraphPad Prims software.

### Cell extracts

To interrogate the levels of TAF1 and ACTB by western blot soluble extracts were made by resuspending cell pellets in RIPA buffer (50mM Tris-Cl pH 7.4, 150mM NaCl, 1% NP40 and 0.25% Na-deoxycholate) supplemented with proteases inhibitors. After 30 minutes in ice lysates were spun down and supernatants collected.

### RNA interference and establishment of stable cell lines

Doxycycline inducible pTRIPZ vectors containing different shRNAs against TAF1 (Table 1) were purchased from Dharmacon. Lentiviral infections were performed as previously described [11]. Stable cell lines were stablished after two weeks of selection with 2μg/ml puromycin and downregulation of TAF1 was assessed by western blot after treatment with 0.5μg/ml doxycycline.

**Table 1.**
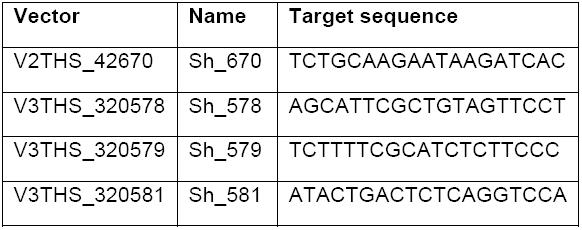
Inducible lentiviral shRNA pTRIPZ vector targeting TAF1 from Dharmacon

### CRISPR-Cas9 gene editing and growth competition assays

gRNAs against TAF1 (Table 2) were designed using the web tool cripsr.mit.edu with a quality score threshold above 80 to minimize off-target effects. Non target gRNAs sequences were previously described [12]. gRNAs against BRD4 were previously described [8]. gRNAs were cloned into pKLV-U6gRNA(BbsI)-PGKpuro2ABFP (Addgene plasmid # 50946). Lentiviral particles were generated as previously described [13] and cells previously modified to express Cas9 using pLentiCas9 Blast (Addgene plasmid # 52962) [14] were infected. Four days post infection, growth competition assays were carried out by mixing an equal number of of BFP+/gRNA expressing cells and non-gRNA transduced parental Cas9 expressing cells (BFP-). The percentage of BFP+ cells was determined by flow cytometry at different days starting the day of the mixing (day 0) and the fold depletion of the percentage of BFP+ cells compared to day 0 was calculated (d0 %BFP+/dN %BFP+). For the statistical analysis the percentage of growth inhibition at day 14 compared to day 0 for each gRNA was calculated and adjusted to the percentage of growth inhibition of the non-target gRNAs. The adjusted percentages of growth inhibition for each gRNA obtained in 3 independent experiments were pooled into categories (NT, non-conserved and conserved amino acids of the bromodomain or the NTK) and the categories were compared using the Tukey-Kramer test [15].

**Table 2.**
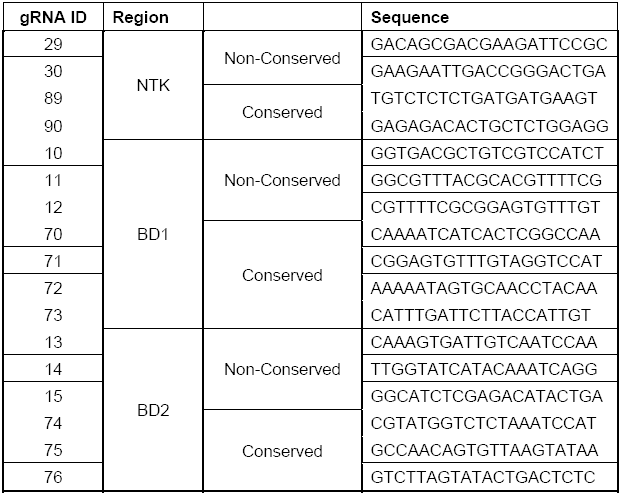
gRNAs for TAF1 used in the CRISPR-Cas9 domain screening:

### RNA-seq

Cells were treated with doxycycline for 10 days or the compounds for 48 hours, collected and total RNA was extracted using the RNeasy kit (Qiagen). Two biological replicates were used per condition. Library construction, sequencing, alignment to human genome hg19 transcript assembly and differential expression were done as previously described [16] using Nextpresso [17]. We compared gene expression in cells treated with compound to vehicle and cells transduced with the TAF1 shRNAs to the non-target shRNA. Genes changing expression with FDR<0.05 were considered as differentially expressed. P-values for the overlap in downregulated or upregulated genes between treatments were calculated using hypergeometric test. Enrichments in biological processes were calculated using PANTHER [18].Transcriptional changes after treatment of K562 cells with JQ1 and CBP30 can be accessed at GEO with accession numbers GSE77295 and GSE110229.

### Gene set enrichment analysis (GSEA)

For GSEAPreranked [19] genes were pre-ranked according to the statistic test of fold change for each treatment obtained in the RNA-seq analysis, setting ‘gene set’ as the permutation method and with 1000 permutations. Gene set SCHUHMACHER_MYC_TARGETS_UP can be found at the Molecular Signatures Database (http://software.broadinstitute.org/gsea/msigdb/index.jsp). Gene set siGATA1_DOWN corresponds to genes downregulated after GATA1 knock down in K562 as previously described [20].

### ChIP-seq analysis

Hg19 aligned bam files for ChIP-seq in K562 were downloaded from ENCODE (GEO accession number GSM803431 for TAF1, GSM733643 for RNA PolII, GSM733656 for H3K27ac and GSM935495 for TBP) and signal densities around the TSS (-/+1kb) of genes were calculated using bamToGFF (https://github.com/bradnerComputation/pipeline/blob/master/bamToGFF.py). Top occupied TSS were selected. P-values for overrepresentation or downrepresentation of genes with top TSS enrichments were determined by chi-square test.

### Cell cycle analysis

Cell pellets were fixed with 70% ethanol in PBS at 4 °C for at least 1 h and stained with propidium iodide (100 ug/ ml) or Hoechst 33342 in the presence of RNase A and 0.1% Triton X-100 at 4 °C for at least 30 min. Cell cycle distribution was measured using a BD LSRFortessa flow cytometer (BD Bio-sciences) and data analyzed using the FlowJo software. Three replicates were used per condition. P-values for differences in percentage of cells in the different phases of the cell cycle were calculated using T-test.

## ACKNOWLEDGEMENTS

This work was funded by Eli Lilly. Authors would like to thank SEPE for support during the writing of this manuscript. We thank M. Serrano and M. Malumbres for fruitful discussions and O. Graña for assistance in the gene expression analysis. We also thank the Flow Cytometry, Bioinformatics and Genomics Units at the CNIO.

Authors declare no conflict of interest.

